# Neurocognitive deficits in controlling aversive memory among insomnia disorders

**DOI:** 10.64898/2026.03.04.709020

**Authors:** Xibo Zuo, Xuanyi Lin, Ziqing Yao, Danni Chen, Jing Liu, Sean Guo, Winny Yue, Ying Yang, Wei Wang, Hongliang Feng, Jihui Zhang, Michael C. Anderson, Shirley Xin Li, Xiaoqing Hu

**Affiliations:** Department of Psychology, The University of Hong Kong, Hong Kong SAR, China; Center for Sleep and Circadian Biology, Northwestern University, Evanston, IL, USA; Department of Neurobiology, Northwestern University, Evanston, IL, USA; Philosophy and Social Science Laboratory of Reading and Development in Children and Adolescents (South China Normal University), Ministry of Education; School of Psychology, South China Normal University, Guangzhou, China; Department of Neurology, The First Affiliated Hospital of Jinan University, Guangzhou, Guangdong, China; Center for Sleep and Circadian Medicine, The Affiliated Brain Hospital, Guangzhou Medical University, Guangzhou, Guangdong, China; Guangdong Engineering Technology Research Center for Translational Medicine of Mental Disorders, Guangzhou, Guangdong, China; Key Laboratory of Sleep and Biological Rhythms of Guangdong Province, Guangzhou, Guangdong, China; Department of Psychiatry, Faculty of Medicine, The Chinese University of Hong Kong, Hong Kong SAR, China; MRC Cognition and Brain Sciences Unit, University of Cambridge, Cambridge, UK; Behavioural and Clinical Neurosciences Unit, University of Cambridge, Cambridge, UK

**Author notes:** Equal contributions (Xibo Zuo, Xuanyi Lin). Corresponding Authors: Shirley X. Li, Xiaoqing Hu, Department of Psychology, The University of Hong Kong, Pokfulam, Hong Kong SAR, China.

**Keywords:** insomnia, emotional memory, Think/No-Think, memory control, EEG

## Abstract

**Background:** Insomnia disorder is a common sleep disturbance characterized by adverse daytime cognitive and emotional impairments, such as repetitive negative thinking and increased psychological distress. Memory control, a key self-regulatory ability to control or inhibit unwanted thoughts and memories, plays an essential role in supporting cognitive functions and emotional well-being. Here, we delineate the neurocognitive mechanisms underlying memory control among individuals with insomnia.

**Methods:** 41 participants meeting DSM-5 criteria for insomnia disorder and 40 healthy sleepers completed an emotional Think/No-Think task, during which participants either retrieved (Think) or suppressed the retrieval (No-Think) of aversive memories in response to memory cues while electroencephalograms were recorded.

**Results:** Linear mixed model analyses with age and depression scores as covariates showed that participants with insomnia exhibited impaired memory control abilities, as evidenced by reduced suppression-induced forgetting in memory recall when compared to healthy sleepers. Electrophysiologically, healthy sleepers showed enhanced right prefrontal theta power in retrieval suppression than in retrieval, indicating elevated needs of inhibitory control during memory control. In sharp contrast, this difference was absent among those with insomnia. Notably, the greater the severity of insomnia symptoms, the smaller the retrieval vs. retrieval suppression theta power differences across participants, linking inefficient top-down control of unwanted memories with low sleep qualities.

**Conclusion:** Individuals with insomnia showed impaired memory control of aversive memories and aberrant electrophysiological activities during retrieval suppression. Future research shall investigate the causal relationship between memory control abilities and insomnia symptoms.

## Introduction

Insomnia is characterized by difficulties in sleep initiation and maintenance, and by early-morning awakening (Riemann et al., 2023). Individuals who experience these sleep symptoms for at least three nights a week, persisting for over 3 months, and accompanied by daytime impairments, meet the diagnosis of insomnia disorder (American Psychiatric Association, 2013). Chronic sleep disruption adversely affects physical health and is associated with an increased risk of various psychiatric disorders, including depression, anxiety, and post-traumatic stress disorder (PTSD) (Baglioni et al., 2011; Hertenstein et al., 2019). To date, insomnia disorder is recognized as the most prevalent sleep disorder, with about 10% of adults meeting its criteria, imposing significant health and socioeconomic burdens (Morin & Buysse, 2024). For example, in China alone, the economic cost of sleep disturbance is estimated to be US$628.15 billion in 2018 due to decreased work productivity (Liu et al., 2025).

Among various cognitive factors that may contribute to insomnia, repetitive negative thinking — persistent, affect-laden intrusive thoughts or “a racing mind” that occur throughout waking and nocturnal periods — plays a crucial role in the etiology and maintenance of insomnia (Espie, 2023; Harvey, 2002, 2005). Recent studies suggest that excessive intrusive thoughts arise not only from “bottom-up” attention biases toward threatening stimuli (Espie, Broomfield, MacMahon, Macphee, & Taylor, 2006) but also from the impairments in “top-down” inhibitory control that would prevent thoughts or memories from intruding into awareness (Harrington & Cairney, 2021). For example, compared to well-rested sleep, a night of sleep deprivation reduced neural activity of the prefrontal cortices, implicating inhibitory control, and increased frequency of memory intrusions (Harrington, Ashton, Sankarasubramanian, Anderson, & Cairney, 2021; Harrington et al., 2025; Zeng, Lau, Li, & Hu, 2021). Moreover, emerging evidence reveals that individuals with insomnia show reduced overnight dissipation and less time-dependent forgetting of aversive memories, suggesting inefficient down-regulation or slowed dissolving of aversive memories during sleep (Cabrera et al., 2024; Wassing, Benjamins, Talamini, Schalkwijk, & Van Someren, 2019; Wassing, Schalkwijk, et al., 2019; Zeng et al., 2025). However, it remains unknown whether people with chronic insomnia, i.e., a clinical condition different from sleep deprivation, would show deficits in controlling unwanted thoughts and memories. Here, we aim to investigate the memory control ability among individuals with chronic insomnia. The ability to control unwanted memories is often studied with the Think/No-Think task (Anderson & Green, 2001), during which participants are confronted with memory reminders that would trigger recollection of the associated memories. In the Think condition, participants are instructed to retrieve and maintain the reminder-associated memory in mind, whereas in the No-Think condition, participants are instructed to stop thinking about the associated memory while paying full attention to the reminder (i.e., retrieval suppression). Such an active stopping of retrieval would reduce intrusions during the Think/No-Think task and further induce forgetting in subsequent memory tests, a phenomenon known as suppression-induced forgetting (Anderson, Crespo-Garcia, & Subbulakshmi, 2025; Anderson & Hulbert, 2021). Retrieval suppression enhances brain activities implicating inhibitory control, such as elevated blood oxygen level dependent (BOLD) signals in the right dorsolateral prefrontal cortex, and increased electrophysiological activities, including the prefrontal 4-8 Hz theta power and event-related potentials N450 (Anderson et al., 2004; Brendan E Depue, Curran, & Banich, 2007; Brendan Eliot Depue et al., 2013; Dutra, Beria, Ligório, & Gauer, 2019; Hu, Bergström, Gagnepain, & Anderson, 2017; Lin et al., 2024). Meanwhile, retrieval suppression down-regulates neural activities implicating episodic retrieval, including reduced BOLD signals in the hippocampus, the medial temporal lobe, and the visual cortex, and reduced electrophysiological activities, such as the parietal P300 (Bergström, de Fockert, & Richardson-Klavehn, 2009; Hanslmayr, Leipold, Pastötter, & Bäuml, 2009; Hellerstedt, Johansson, & Anderson, 2016; Hu, Bergström, Bodenhausen, & Rosenfeld, 2015; Mecklinger, Parra, & Waldhauser, 2009).

Here, we investigated the neurocognitive mechanism underlying memory control among individuals with insomnia disorder. We hypothesized that individuals diagnosed with insomnia disorder would exhibit a deficit of memory control, indicated by reduced suppression-induced forgetting and electrophysiological signals related to retrieval suppression (e.g., prefrontal theta power). Elucidating the mechanisms of memory control could provide not only the mechanistic insights into why people with insomnia tend to have uncontrollable repetitive negative thinking, but also inform the development of cognitive interventions that may alleviate insomnia symptoms.

## Method

### Participants

We recruited 48 participants with insomnia via mass emails, social media, posters/ flyers across university campuses and local communities. Six were excluded due to non-compliance with task instructions, and one due to excessive EEG artifacts (> 50%), yielding 41 participants with insomnia disorder and 40 healthy sleepers (data from Lin et al., 2024). Sample sizes (40 per group) aligned with recent emotional Think/No-Think studies (Küpper, Benoit, Dalgleish, & Anderson, 2014; Lin et al., 2024; Meyer & Benoit, 2023).

Participants were pre-screened to ensure their eligibility. All participants were aged 18-35 years. Healthy sleepers had no current or past history of insomnia. Participants with insomnia met 1) the diagnostic criteria of insomnia disorder in the 5^th^ edition of the Diagnostic and Statistical Manual of Mental Disorders (DSM-5) (American Psychiatric Association, 2013), and 2) scored ≥ 10 on the Insomnia Severity Index (ISI), aligning with clinical thresholds for the community samples (Morin, Belleville, Bélanger, & Ivers, 2011). Exclusion criteria for both groups included: 1) night shift worker; 2) current neurological/psychiatric diagnoses (other than insomnia in the insomnia group); and 3) current prescription medication use or medical conditions influencing cognitive functions or sleep. All participants had normal or corrected-to-normal vision and hearing. This study was approved by the Human Research Ethics Committee of the University of Hong Kong (Ref. EA210341). Participants provided their written informed consent before participation and received full debriefing and monetary compensation (HKD$250 = US$32) upon completion.

### Procedure

Potential interested participants completed online sign-ups followed by attending a screening clinical interview administered by trained interviewers using 1) the Diagnostic Interview for Sleep Patterns and Disorder-Chinese (DISP; Merikangas et al., 2014), which assesses major sleep disorders (e.g., hypersomnia) based on the DSM-5 and the International Classification of Sleep Disorders (ICSD) criteria (American Psychiatric Association, 2013; Sateia, 2014), and 2) the Mini-International Neuropsychiatric Interview (MINI; Sheehan et al., 1998) to assess major neuropsychiatric disorders (e.g., major depressive disorder). Eligible participants were subsequently invited to complete EEG testing within seven days after the clinical interview. Upon arrival, participants completed self-report questionnaires assessing insomnia severity (ISI; Morin et al., 2011), depressive symptoms (Beck Depression Inventory-LJ, BDI-II; Beck, Steer, & Brown, 1996), and anxiety (State-Trait Anxiety Inventory, STAI; Spielberger, Gonzalez-Reigosa, Martinez-Urrutia, Natalicio, & Natalicio, 1971). Questionnaire details are provided in the Supplementary Materials.

Participants then completed the emotional Think/No-Think task, comprising: 1) encoding; 2) Think/No-Think with EEG recording; 3) cued recall (Figure 1a). An implicit affect task was administered before and after cued recall to assess the aftereffect of retrieval suppression on spontaneous affective judgments. Procedure and results about this task are reported in the Supplementary Materials. Finally, participants completed 3 self-report questionnaires to verify task compliance and strategy utilization. Tasks were programmed by E-Prime® 2.0 (Psychology Software Tools, Inc., Pittsburgh, USA). All task procedures were consistent with those of a previous study (Lin et al., 2024).

**Figure 1.**
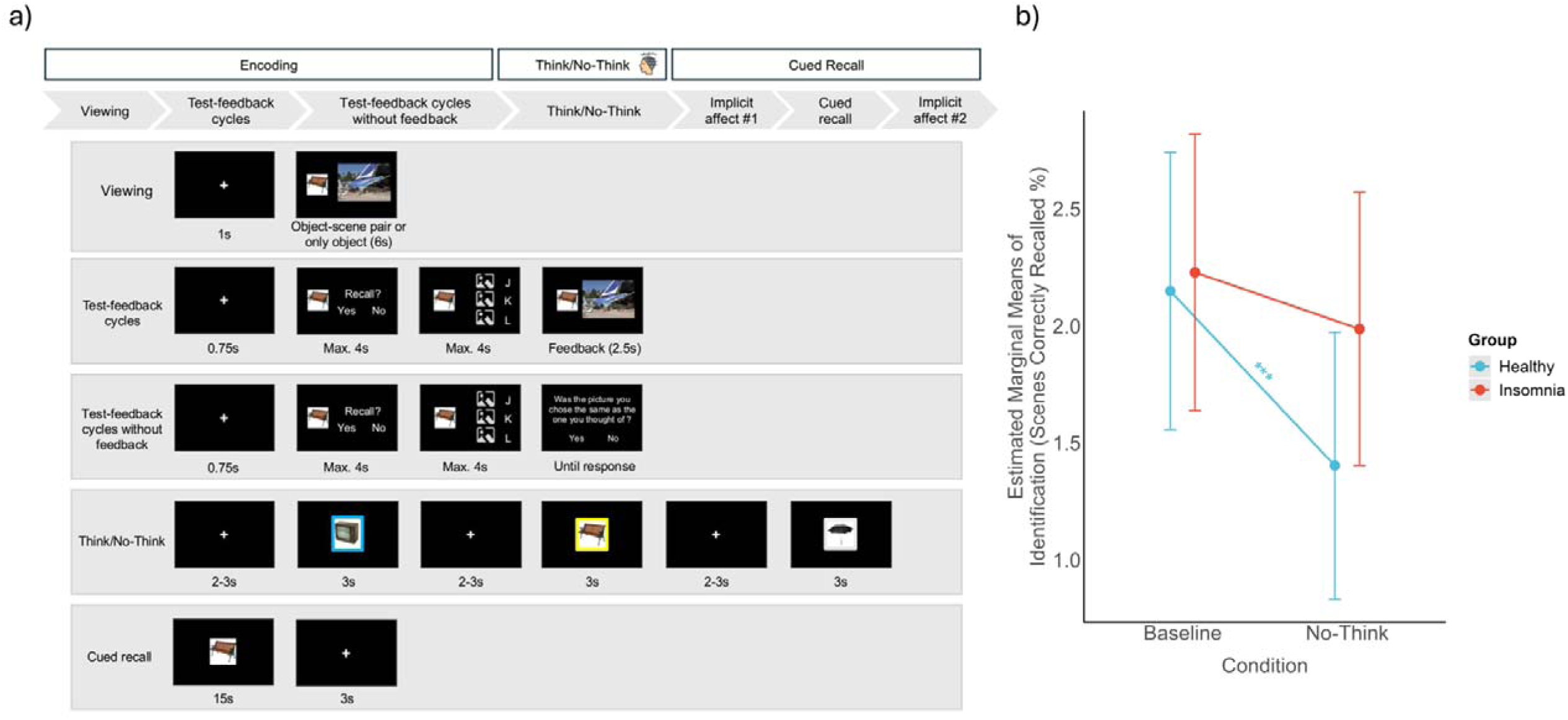
a) Experimental design and task details. The emotional Think/No-Think task included 3 phases: encoding, Think/No-Think, and cued recall phases. The stimuli were selected from validated sets in accordance with copyright regulations (Küpper et al., 2014). b) Memory performance of Identification in the cued recall task: healthy sleepers showed worse memories for the No-Think condition compared to the Baseline condition, suggesting a suppress-induced forgetting effect, whereas individuals with insomnia did not. Error bars represent 95% confidence intervals (CIs). Asterisks present a significant difference between conditions (\**p* < .05; \*\**p* < .01; \*\*\**p* < .001).

### Emotional Think/No-Think Task

#### Encoding

Encoding comprised viewing and test-feedback cycles. Stimuli included 42 object-aversive scene picture pairs and 6 unpaired objects, selected from validated sets (Küpper et al., 2014). During viewing, each pair or unpaired object was presented for 6s with 1s inter-trial interval (ITI) in randomized order. For object-scene pairs, participants memorized the association between the left-sided object and the right-sided scene and focused on details in the scene. For unpaired objects, participants simply viewed the object. In the subsequent test-feedback cycles, where unpaired objects were not shown, participants were shown an object and indicated within 4s whether they could recall the associated scene. For ‘yes’ responses, three scenes from the encoding phase were presented vertically, and participants selected the correct scene within 4s. Feedback showing the correct object-scene pair was presented for 2.5s, regardless of accuracy. The test-feedback cycles were repeated until participants reached 60% accuracy. All participants reached this accuracy criterion within three test-feedback cycles. Once reaching this criterion, a final test cycle without feedback was performed.

#### Think/No-Think

This phase included 24 previously objects from the encoded object-scene pairs and 6 unpaired objects. Trial began with a fixation cross with 2–3s in 100ms steps (i.e., 2s, 2.1s, …, 3s), followed by an object presented in either yellow, blue, or white color frames for 3s. Yellow and blue frames indicated the Think or No-Think condition, respectively, with the color-condition assignment counterbalanced across participants. White frames indicated the Perceptual Baseline condition. Frame colors for objects were consistent across all trials. Participants were instructed to fixate their eyes on the object image for all conditions. In the Think condition, participants were instructed to actively retrieve and maintain the associated scene in mind throughout the presentation of the object. In the No-Think condition, participants were instructed to focus on the object while preventing any associated memories from coming into mind. If any scene-associated memories came to mind, they need to expunge intruding memories from their mind. In the Perceptual Baseline condition, participants sustain attended to the objects during the whole trial without any further instructions. The 24 previously encoded paired objects were assigned to Think and No-Think conditions (12 each). The 6 unpaired objects were assigned to the Perceptual Baseline condition. Each object was presented 10 times (300 trials in total). The remaining 12 previously encoded paired objects served as the Baseline condition and were not presented in this phase. 6 additional paired objects were used for practice.

#### Cued recall

Participants were presented with the 36 paired objects (12 Think, 12 No-Think, 12 Baseline) in random order without color frames. For each object, participants verbally described the associated scene within 15s. Verbal responses were audio-recorded for later scoring. Trials were separated by a 3s ITI. Unpaired objects from the Perceptual Baseline condition were shown. Assignment of 12 object-scene pairs to Think, No-Think, and Baseline conditions was counterbalanced across participants, ensuring comparable materials. Valence and arousal level for each object-scene set were comparable (see Lin et al., 2024).

### Electrophysiological acquisition, processing, and analysis

Electroencephalography (EEG) was continuously recorded during the Think/No-Think phase via a 64-channel cap based on the international 10-5 system (eego mylab, ANT Neuro, Germany). EEG data sampled at 500 Hz, referenced online to CPz, with AFz as ground. Electrooculogram (EOG) was recorded below the left orbit, and impedances were maintained below 20 kΩ.

Preprocessing was conducted in Matlab using EEGLAB (Delorme & Makeig, 2004) and ERPLAB (Lopez-Calderon & Luck, 2014). Data were downsampled to 250 Hz, band-pass filtered (0.1-60 Hz), notch-filtered at 50 Hz, re-referenced to the common average, and cleaned via Independent Component Analyses (ICA) to remove ocular and muscle artifacts. Trials exceeding ±100 μV were rejected. The number of trials did not differ across groups, conditions, or object-scene sets (*Fs* < 3.29, *ps* > .074).

For event-related potential (ERP) analyses, epochs were segmented from -200 to 1500ms relative to the cue onset, using -200 to 0ms for baseline correction. Mean amplitudes were quantified for two components: the fronto-central N450 (FC1, FC2, C1, C2, FCz, Cz) from 300-500ms and the parietal P300 (Pz, CP1, CP2, P1, P2) from 300-800ms (Bergström et al., 2009; Chen et al., 2012; Hellerstedt et al., 2016; Hu et al., 2015; Lin et al., 2024).

Time-frequency decomposition was performed in Fieldtrip (Oostenveld, Fries, Maris, & Schoffelen, 2011). To mitigate edge artifacts, epochs were trimmed to -500 to 3000 ms relative to cue onset. Morlet wavelet transforms were applied with cycle parameters increasing linearly from 3 at 2.8 Hz to 15 cycles at 30 Hz, across 22 logarithmically spaced frequency bins. Spectral power was normalized by z-score transformation of all trials relative to the baseline period (-500 to -200 ms). Analyses focused on the theta (4-8 Hz) power from 200 to 400 ms over right prefrontal electrodes (AF4, F4, F6) (Lin et al., 2024), consistent with the established role of the right dorsolateral prefrontal cortex (rDLPFC) in mnemonic inhibition (Anderson & Hulbert, 2021).

### Statistical analysis

Analyses were conducted in R version 4.5.1 (R Core Team, 2025). Group differences in demographic and self-reported measures were assessed using chi-square tests for categorical variables and independent t-tests for continuous variables.

Memory performance was quantified using three standard indices in the emotional Think/No-Think research: Identification, Gist, and Detail. Two trained raters, blind to condition and group, scored all responses independently. Identification was dichotomous (1 = sufficient detail to uniquely identify the scene; 0 = insufficient), with rater disagreements assigned 0.5. Gist was the ratio of correctly reported critical thematic elements of each scene to predefined total elements (2 to 4 per scene). Detail was the number of accurate, scene-specific descriptive units. Interrater agreement was high: Identification: *r* = 0.84; Gist: *r* = 0.93; Detail: *r* = 0.93).

Linear mixed-effects model (LMM) and cumulative link mixed models (CLMM) were fitted to item-level data to account for participant and item variability via the lme4 and the ordinal package (Bates, Mächler, Bolker, & Walker, 2015; Christensen, 2018). Treatment coding was applied. Tukey-adjusted post-hoc pairwise comparisons were computed using the emmeans package (Lenth, 2022). We reported the marginal R^2^ (variance explained by fixed factors only) and the conditional R^2^ (variance explained by both fixed and random factors) for LMM, and the McFadden’s pseudo-R^2^ for CLMM to indicate the goodness-of-fit of our models (S. S. Mangiafico, 2016; Nakagawa & Schielzeth, 2013), by the rcompanion and the MuMIn package (Burnham & Anderson, 2002; S. Mangiafico & Mangiafico, 2017). Odds ratio (OR) and Cohen’s *d* with 95% Confidence Interval (CI) were obtained using the effectsize package for any pairwise comparisons (Ben-Shachar, Lüdecke, & Makowski, 2020). LMM was applied to the Gist and Detail; Identification was analyzed via CLMM due to its ordinal scoring (0, 0.5, 1). Suppression-induced forgetting was assessed with models including fixed effects of Group (healthy sleeper vs. insomnia), Condition (No-Think vs. Baseline), their interaction, and covariates age and depression, with random intercepts for participants and items.

Identification or Gist or Detail ∼ 1 + Group * Condition + age + depression + (1 | subject) + (1 | item)

Facilitation (i.e., enhanced memory for the Think condition compared with the Baseline condition) used the same model and was reported in the Supplementary Materials. EEG data were analyzed using analogous LMMs, except that Condition included only Think and No-Think conditions due to fewer Perceptual Baseline trials.

N450 or P300 or theta ∼ 1 + Group * Condition + age + depression + (1 | subject) + (1 | item)

Spearman’s rho was computed to examine relationships between insomnia severity (ISI scores), suppression-induced forgetting, and EEG measures. suppression-induced forgetting was calculated by subtracting memory performance in the No-Think condition from that in the Baseline condition. To control for item-level differences in memorability and intrusiveness across counterbalancing pair sets, ISI scores, suppression-induced forgetting, and EEG measures were z-normalized within participants’ counterbalancing object-scene pair sets (Nardo & Anderson, 2024). Semipartial correlations covaried out the confounding effect of depressive symptomatology. Statistical significance was defined as *p* < .05 threshold (two-tailed).

## Results

### Sample characteristics

Participants with insomnia had significantly higher levels of insomnia and depressive symptoms, and were older than healthy sleepers (*p*s < .001). Age and depression scores were subsequently included as covariates in follow-up analyses to account for potential confounding effects. No significant differences emerged between two groups across other demographical and self-report variables (*p*s > .05; see Table 1). Moreover, encoding accuracy in the final test-feedback cycle was comparable across groups of participants, conditions, and object-scene pair sets (*Fs* < 1.93*, ps > .*169, *η^2^_p_* < 0.05).

**Table 1.**
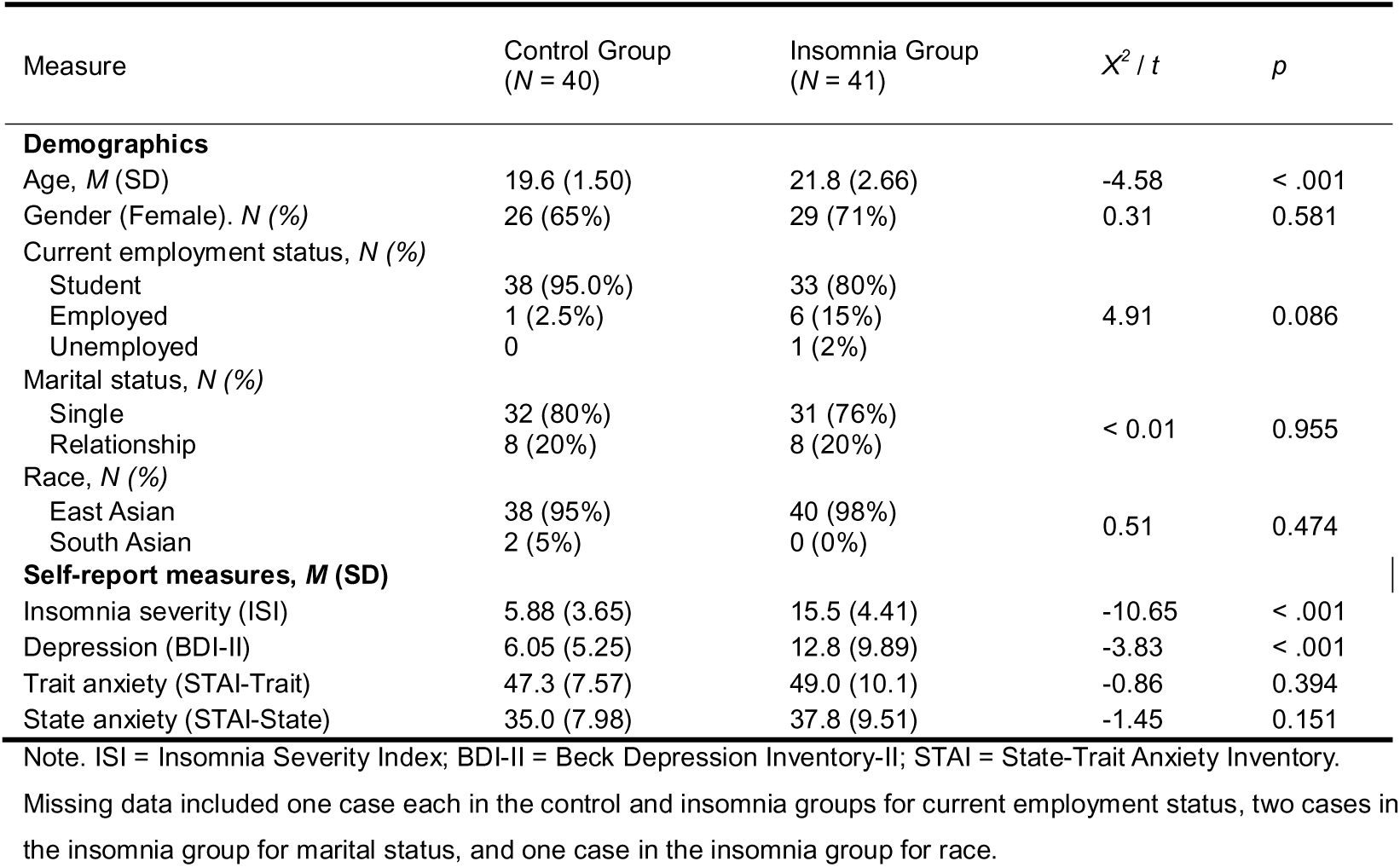
Self-report characteristic differences between the healthy sleeper group and the insomnia group (N = 81).

| Measure | Control Group<br>(N = 40) | Insomnia Group<br>(N = 41) | $\chi^2 / t$ | *p* |
| --- | --- | --- | --- | --- |
| **Demographics** |  |  |  |  |
| Age, *M* (SD) | 19.6 (1.50) | 21.8 (2.66) | -4.58 | < .001 |
| Gender (Female). *N* (%) | 26 (65%) | 29 (71%) | 0.31 | 0.581 |
| Current employment status, *N* (%) |  |  |  |  |
| Student | 38 (95.0%) | 33 (80%) |  |  |
| Employed | 1 (2.5%) | 6 (15%) | 4.91 | 0.086 |
| Unemployed | 0 | 1 (2%) |  |  |
| Marital status, *N* (%) |  |  |  |  |
| Single | 32 (80%) | 31 (76%) |  |  |
| Relationship | 8 (20%) | 8 (20%) | < 0.01 | 0.955 |
| Race, *N* (%) |  |  |  |  |
| East Asian | 38 (95%) | 40 (98%) |  |  |
| South Asian | 2 (5%) | 0 (0%) | 0.51 | 0.474 |
| **Self-report measures, *M* (SD)** |  |  |  |  |
| Insomnia severity (ISI) | 5.88 (3.65) | 15.5 (4.41) | -10.65 | < .001 |
| Depression (BDI-II) | 6.05 (5.25) | 12.8 (9.89) | -3.83 | < .001 |
| Trait anxiety (STAI-Trait) | 47.3 (7.57) | 49.0 (10.1) | -0.86 | 0.394 |
| State anxiety (STAI-State) | 35.0 (7.98) | 37.8 (9.51) | -1.45 | 0.151 |

Note. ISI = Insomnia Severity Index; BDI-II = Beck Depression Inventory-II; STAI = State-Trait Anxiety Inventory.
Missing data included one case each in the control and insomnia groups for current employment status, two cases in the insomnia group for marital status, and one case in the insomnia group for race.

### Suppression-induced forgetting

Consistent with our hypothesis, participants with insomnia exhibited reduced suppression-induced forgetting effects in memory identification when compared to healthy sleepers (for descriptives, see Table 2). For Identification, we observed a significant Condition by Group interaction (χ*^2^*(1) = 3.97, *p* = .046; see Figure 1b): Healthy sleepers showed significant below-baseline forgetting (No-Think < Baseline, *M* = -9.7 %, *z* = -4.11, *p* < .001, OR = 0.47, 95% CI [0.33, 0.68]). In contrast, participants with insomnia showed no significant difference between No-think and Baseline conditions (*M* = -3.4 %, *z* = -1.37, *p* = .172, OR = 0.79, 95% CI [0.55, 1.11]). We also found a significant Condition effect (χ*^2^*(1) = 15.20, *p* < .001) indicating the suppression-induced forgetting, and a non-significant Group effect (χ*^2^*(1) = 1.49, *p* = .223). The McFadden’s pseudo-R² for this model was low (pseudo-R² = 0.01).

**Table 2.** Memory performance in the cued recall phase for the healthy sleeper group and the insomnia group (N = 80).

| Measures<br>*M (SD)* | Control Group (*N* = 40) |  |  |  |  | Insomnia Group (*N* = 40) |  |  |  |  |
| --- | --- | --- | --- | --- | --- | --- | --- | --- | --- | --- |
|  | No-Think | Baseline | Think | Suppression-Induced Forgetting | Facilitation | No-Think | Baseline | Think | Suppression-Induced<br>Forgetting | Facilitation |
| **Identification** |  |  |  |  |  |  |  |  |  |  |
| Scenes Correctly<br>Recalled (%) | 0.74 (0.19) | 0.83 (0.13) | 0.86 (0.12) | 0.09 (0.14) | 0.02 (0.12) | 0.79 (0.17) | 0.82 (0.13) | 0.84 (0.13) | 0.03 (0.14) | 0.02 (0.12) |
| **Gist** |  |  |  |  |  |  |  |  |  |  |
| Scenes Correctly<br>Recalled (%) | 0.58 (0.18) | 0.61 (0.13) | 0.66 (0.11) | 0.03 (0.13) | 0.05 (0.14) | 0.57 (0.16) | 0.60 (0.14) | 0.60 (0.16) | 0.02 (0.12) | < -0.01 (0.12) |
| **Detail** |  |  |  |  |  |  |  |  |  |  |
| Number of correct<br>details recalled (%) | 7.23 (1.94) | 7.67 (1.35) | 8.24 (1.78) | 0.44 (1.27) | 0.57 (1.30) | 7.13 (2.07) | 7.69 (1.86) | 7.86 (1.98) | 0.54 (1.44) | 0.16 (1.30) |

Note. Suppression-Induced Forgetting = Baseline – No-Think. Facilitation = Think – Baseline. We missed one participant in the insomnia group because the recording files were broken.

No significant Condition by Group differences were found on Gist (*F*(1, 1804.37) = 0.03, *p* = .874) and on Detail (*F*(1, 1804.06) = 0.26, *p* = .609) (for detailed results, see Supplementary Materials).

### Neural processes during retrieval suppression

Impaired suppression-induced forgetting among participants with insomnia suggested impaired top-down inhibitory control of aversive memories. Complementing the behavioral suppression-induced forgetting, we found a significant Condition by Group interaction for the right prefrontal theta power (*F*(1, 1830.54) = 6.30, *p* = .012; see Figure 2): Healthy sleepers exhibited enhanced theta power for No-Think condition compared to Think condition (*t*(1831) = 3.51, *p* = .001, Cohen’s *d* = 0.23, 95% CI [0.10, 0.36]), whereas the insomnia group showed no significant difference between No-Think and Think conditions (*t*(1827) = -0.02, *p* = .988, Cohen’s *d* < -0.01, 95% CI [-0.13, 0.13]). We observed a significant Condition effect (*F*(1, 1828.21) = 6.20, *p* = .013) and non-significant Group effect (*F*(1, 76.95) = 3.40, *p* = .069). The marginal and conditional R^2^ of our model were low (0.04 and 0.35, respectively), indicating that the fixed effects accounted for only a small proportion of the variance (4%), whereas a larger portion (35%) was explained when random effects were included.

**Figure 2.**
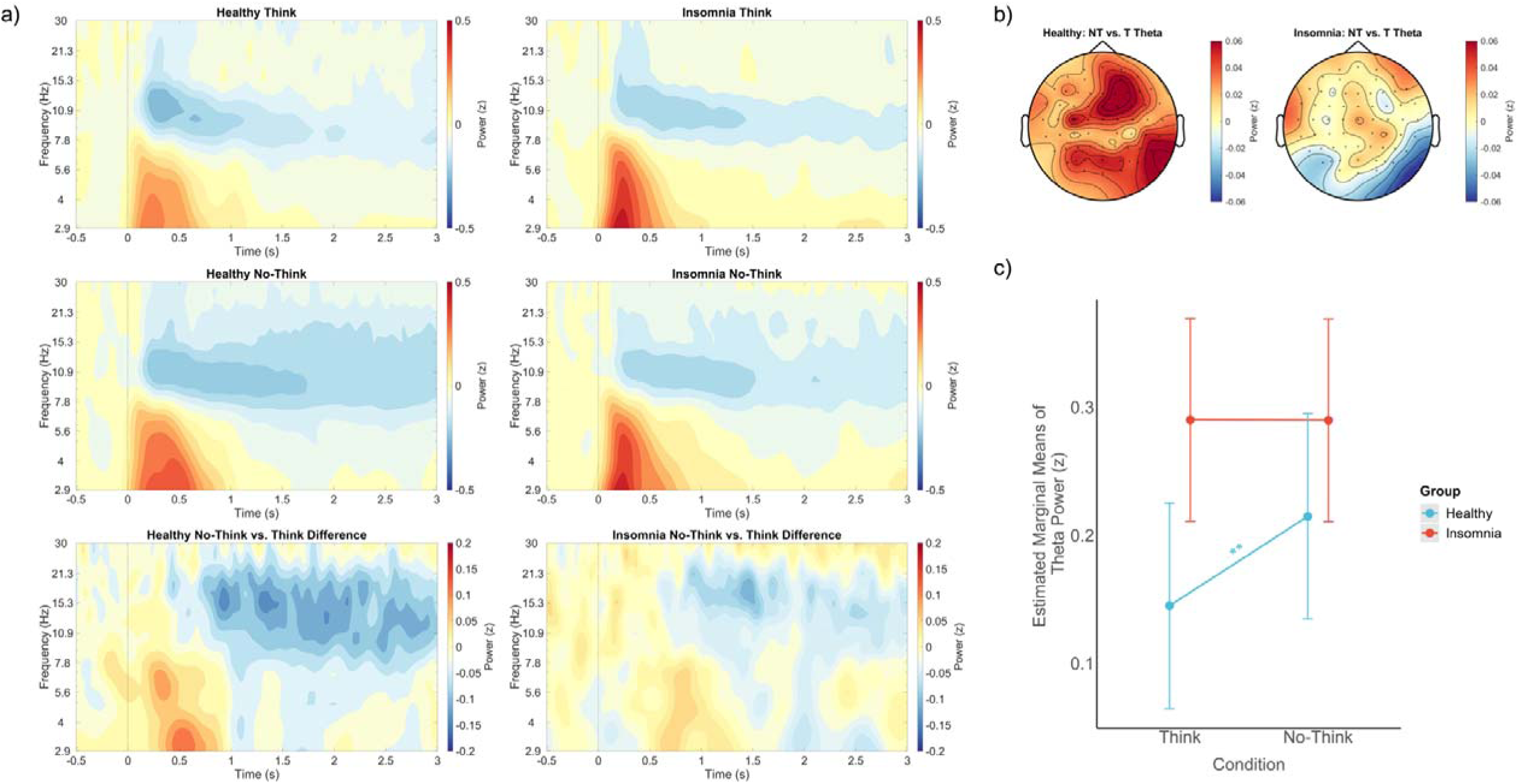
a) The time-frequency plots of Think, No-Think conditions and condition differences (subtracting the Think condition from the No-Think condition) in healthy sleepers and the insomnia group at the right prefrontal area (AF4, F4, F6). b) Topographies of theta oscillations (4-8 Hz) difference averaged on 200-400ms between the No-Think and the Think conditions. NT= No-Think; T = Think. c) Theta power during the Think/No-Think phase: healthy sleepers exhibited enhanced right prefrontal theta power during the No-Think condition compared to the Think condition, whereas individuals with insomnia showed no such difference. Asterisks present significant difference between conditions (\**p* < .05; \*\**p* < .01; \*\*\**p* < .001).

For N450 and P300, we did not observe a significant Condition by Group interaction (N450: *F*(1,1828.13) = 0.01, *p* = .920; P300: *F*(1, 1829.45) = 2.01, *p* = .157; detailed results see Supplementary Materials).

### Correlations between ISI, memory performance, and neural activities

We first normalized all measures within counterbalancing conditions across all participants. A robust negative correlation was observed between the difference in right prefrontal theta power differences (No-Think vs. Think conditions) and ISI scores (rho = -0.29, *p* = .009; see Figure 3): higher insomnia severity was associated with reduced theta power in the No-Think relative to the Think condition. Importantly, this correlation remained significant after controlling for depressive symptoms using semipartial correlation analyses (adjusted rho = -0.29, *p* = .009). However, neither right prefrontal theta nor ISI scores demonstrated a significant correlation with the suppression-induced forgetting measures (*p*s > .141). Next, correlation analyses were conducted separately within each group. We observed the same significant negative correlation between the difference in right prefrontal theta power (No-Think vs. Think conditions) and ISI scores among the healthy group (rho = -0.37, *p* = .018). This association was significant even after controlling for the depressive symptoms (adjusted rho = -0.35, *p* = .028). In contrast, no significant correlation between the difference in right prefrontal theta power (No-Think vs. Think conditions) and ISI scores was found in the insomnia group (rho = -0.10, *p* = .533).

**Figure 3.**
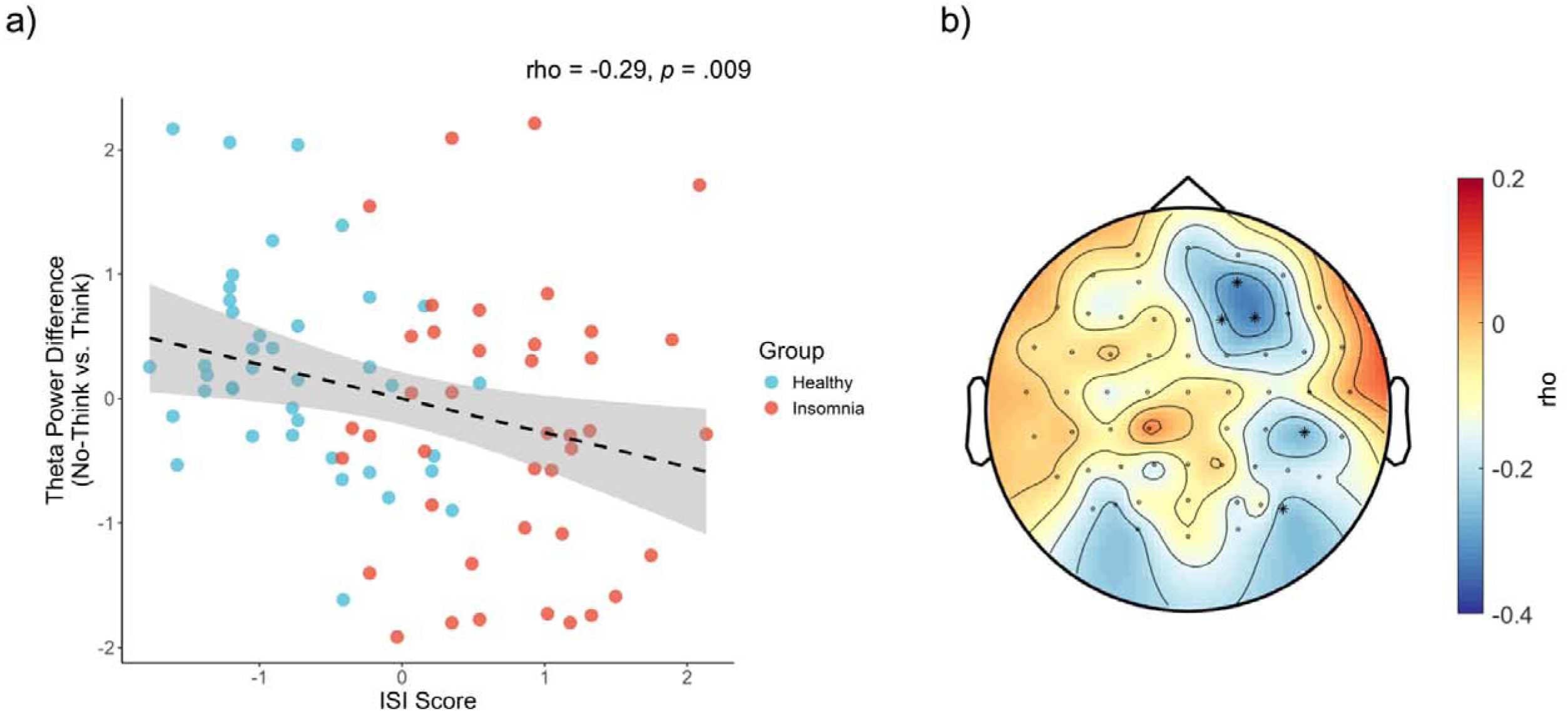
a) Scatter plots depicted the correlation between z-normalized insomnia severity and theta power difference between No-Think and Think conditions (subtracting the Think condition from the No-Think condition) at the right prefrontal (AF4, F4, F6) area among all samples; b) Topographical distribution of the correlation between z-normalized insomnia severity and theta power difference (No-Think vs. Think) at each electrode. Electrodes with significant correlations with insomnia severity are highlighted (uncorrected, αs = 0.05).

## Discussion

The ability to voluntarily control unwanted thoughts and memories is instrumental for cognitive function and for mental well-being. Here, we found that participants with insomnia showed compromised memory control on negative memories, as evidenced by diminished suppression-induced forgetting effect and reduced suppression-related theta power difference when they were supposed to stop the retrieval of memories, compared to healthy sleepers. Moreover, theta power differences are associated with the severity of insomnia symptoms across participants. Together, our study provides novel evidence suggesting that chronic sleep disruption is associated with deficits in controlling aversive memories and thoughts.

Growing evidence reveals that people with insomnia show impaired inhibitory control abilities across domains, including motor inhibition (e.g., prolonged reaction times in stop-signal tasks) and emotional regulation (Ding et al., 2023; Kim et al., 2025; Ling et al., 2021; Zhao et al., 2018). We demonstrate that insomnia also shows inefficient control of aversive memories, suggesting that the insomnia disorder can be characterized by a domain-general deficit of inhibitory control, impairing the top-down control of motor, emotion, and memory responses. The domain-general hypothesis of prefrontal cortex-mediated inhibition posits that shared neuroanatomical and neurophysiological mechanisms govern the voluntary control of motor, emotional, and mnemonic processes (Brendan E Depue, Orr, Smolker, Naaz, & Banich, 2016; Wessel & Anderson, 2024). Notably, deficits in memory control are also observed in other psychiatric disorders that are frequently comorbid with insomnia, including depression, anxiety, and PTSD (Catarino, Küpper, Werner-Seidler, Dalgleish, & Anderson, 2015; Marzi, Regina, & Righi, 2014; Zhang, Xie, Liu, & Luo, 2016; for a meta-analysis, see Stramaccia, Meyer, Rischer, Fawcett, & Benoit, 2021). These converging findings highlight the critical role of high-quality sleep in supporting efficient top-down memory control and in safeguarding mental well-being.

Consistent with recent meta-analyses reporting an aggregated suppression-induced forgetting effect (*d*LJ≍LJ0.20–0.40) (Clark et al., 2025; Stramaccia et al., 2021), healthy sleepers in our study demonstrated a significant suppression-induced forgetting effect for Identification (*d* = 0.67) and Details (*d* = 0.37). In contrast, participants with insomnia demonstrated impaired forgetting of aversive memories on Identification (*d* = 0.25), consistent with the suppression-induced forgetting effect size observed among (sub)clinical individuals with anxiety, depression (*d* = 0.17, 95% CI [-0.09, 0.43]) (Stramaccia, Meyer, Rischer, Fawcett, & Benoit, 2021). This result aligns with poor thought control ability reported in insomnia, particularly for uncontrollable affect-laden thoughts that occur throughout waking and nocturnal periods (Harvey, 2003; Lemyre, Belzile, Landry, Bastien, & Beaudoin, 2020; Schmidt, Harvey, & Van der Linden, 2011). Importantly, both the insomnia and the healthy sleeper groups demonstrated comparable performance during the encoding phase, suggesting that the deficits of memory control could not be attributed to deficits in initial memory encoding. Corroborating this account, participants with insomnia (*M* < -0.2%) also exhibited significantly reduced facilitation for gist relative to healthy sleepers (*M* = 4.7%) (for detailed results, see Supplementary Materials). Thus, reduced suppression-induced forgetting in Identification among participants with insomnia is best explained by the deficits in inhibitory control, but not due to insufficient encoding, nor to biases in retrieving negative memories.

While we observed impaired forgetting of memories in Identification among individuals with insomnia relative to healthy sleepers, both healthy and insomnia sleepers showed significant suppression-induced forgetting in memory details. Insomnia sleepers thus show an intriguing pattern of memory control: retrieval suppression reduced details but not memory identification. A potential explanation could be that insomnia sleepers may also show overgeneral memory that is often documented among individuals with psychiatric disorders, particularly those with depression and PTSD (Hallford, Rusanov, Yeow, & Barry, 2021; Ono, Devilly, & Shum, 2016). Overgeneral memory is characterized by a tendency to recall overgeneralized autobiographical memories (i.e., Identification) with reduced specificity (i.e., Details, Sumner, Griffith, & Mineka, 2010). Based on the CaRFAX model (Williams, 2006), overgeneralized memory is affected by 1) rumination about distressing general autobiographical themes, 2) functional avoidance of the risk to experience affective disturbance, and 3) impairment in executive control of retrieving specific memories. In the context of insomnia, when confronted with negative memory reminders, they may recall fewer details either to avoid the risk of affective disturbance or due to executive dysfunction. This tendency may contribute to the observed forgetting effect for memory details. Supporting this idea, previous studies found that shorter sleep duration and sleep deprivation are correlated with reduced specificity of autobiographical memories (Barry, Takano, Boddez, & Raes, 2019; Zare Khormizi, Salehinejad, Nitsche, & Nejati, 2019). Based on the current results, future research should investigate the relationship between insomnia and overgeneralized autobiographical memories.

Electrophysiological analyses focused on early engagement of inhibitory control as indicated by the right prefrontal theta power within 0-500 ms post-cue window. We found that compared to healthy sleepers, insomnia sleepers showed deficits in the early inhibitory control, as evidenced by the non-significant theta power differences between retrieval and retrieval suppression conditions. The prefrontal theta activity is a well-established neural signature of inhibitory control, and has been repeatedly found in previous Think/No-Think research (Crespo-García, Wang, Jiang, Anderson, & Lei, 2022; Brendan Eliot Depue et al., 2013; Lin et al., 2024). Here, reduced early theta activity suggests reduced or inefficient recruitment of the prefrontal cortex during retrieval suppression among insomnia sleepers. Furthermore, across participants, the severity of insomnia symptoms was associated with diminished suppression-related theta power differences, even after controlling for depressive symptoms. Thus, beyond the group-level insomnia vs. healthy sleeper comparisons, we provide direct evidence linking insomnia symptoms and neural activity implicating inhibitory control across individuals.

Our results bear clinical implications for developing interventions targeting the control of aversive or unwanted thoughts/memories (Baker, Baldwin, & Garner, 2015; Ballesio, Ottaviani, & Lombardo, 2019). Recent research demonstrates that a 3-day cognitive training in suppressing future worries and fear would reduce vividness and emotional salience of fears, and improve overall mental health outcomes without inducing a paradoxical rebound effect (Mamat & Anderson, 2023). This cognitive training on confronting reminders of negative personal events or fearful thoughts, and subsequently suppressing associated memories integrates two mechanisms: (1) active forgetting of distressing imagery (Benoit, Davies, & Anderson, 2016; Phelps & Hofmann, 2019) and (2) controlled engagement of extinction circuitry (Anderson & Floresco, 2022; Beckers et al., 2023; Craske, Sandman, & Stein, 2022). This combined approach is theorized to regulate emotional response to threatening information and distressful memories. Based on this research and our results, one possible intervention is to train people with insomnia to suppress aversive memories and sleep-related worries, which may help to down-regulate emotional responses to sleep-related threats and worries, and thus improve sleep quality. Furthermore, future studies could also investigate whether Cognitive-Behavioral Therapy for Insomnia (CBT-I), the current gold-standard non-pharmacological treatment for insomnia, could enhance the ability to control aversive memories by improving sleep quality and insomnia symptoms.

Several limitations should be considered. First, given the limited spatial resolution of scalp EEG, to what extent insomnia severity might modulate the functional connectivity implicating memory control remained unclear. Future research shall employ fMRI to examine the precise neuroanatomical and functional network underlying memory suppression among insomnia sleepers. Second, the cross-sectional design precluded causal inferences regarding the relationship between impaired retrieval suppression and insomnia symptoms. Although greater insomnia severity was associated with reduced electrophysiological activities reflecting inhibitory control in healthy sleepers, longitudinal investigations are required to disentangle whether pre-existing inhibitory control deficits predispose people to insomnia development or whether chronic sleep disruption degrades the function of inhibitory control.

In conclusion, the present study provides insights regarding the neurocognitive mechanisms underlying failures of mnemonic control in insomnia. Future studies with longitudinal designs are needed to further investigate the causal relationship between thought control deficits and insomnia severity. Interventions that improve memory control abilities and sleep quality in this population can help establish the causal relationship between memory control abilities and insomnia symptoms, and eventually safeguard mental health.

## Data availability statement

The data and software/scripts used in this study are available at https://osf.io/3adw2/.

## Acknowledgements

The authors gratefully acknowledge the valuable contributions of all participants and the people involved in recruitment and assessments: Jiefan Ling, Hao Fong Sit, Chi Lok Enoch Wong, and Chan Chi Lam.

## Ethical standards

The authors assert that all procedures contributing to this work comply with the ethical standards of the relevant national and institutional committees on human experimentation and with the Helsinki Declaration of 1975, as revised in 2008.

