## Supplemental Materials for "Neurocognitive deficits in controlling aversive memory among insomnia disorders"

Supplementary Materials

### Method

#### Emotional Think/No-Think Task

*Encoding*. For the test-feedback cycles, all participants reached the accuracy criterion (60%) within three test-feedback cycles: 57 participants succeeded in the first cycle, 23 participants succeeded in two cycles, and one participant used three cycles.

*Think/No-Think.* To assess compliance with the instructions during the No-Think condition, participants completed three self-report items following the Think/No-Think phase. Each item was rated on a five-point scale (1 = Never, 2 = Rarely, 3 = Sometimes, 4 = Often, 5 = Always). The items were: 1) When I saw a yellow/blue framed object, I quickly checked to see if I remembered the associated scene; 2) After a yellow/blue framed object went off the screen, I checked to see if I still remembered the associated scene; 3) When I saw a yellow/blue framed object, I thought about the scene that went with it to improve my memory for that object-scene pair. A total compliance score was computed by summing responses across the three items. Participants whose total score was less than 10 were classified as non-compliant with the No-Think instruction.

#### Implicit affect task

During this task, thirty-six objects from the object-scene pairs (12 Think, 12 No-Think,12 Baseline) from the encoding phase in the emotional Think/No-Think task were presented. Each object was presented four times, yielding a total of 144 trials. Each trial began with a central fixation cross for 0.5s, followed by an object for 0.075s. A blank screen then appeared for 0.125s, after which a meaningless character was shown for 0.1s before being masked. Participant was instructed to perform an affective judgement towards the character, categorizing it as either positive or negative by pressing the far-left key via left index finger or the far-right key via right index finger, respectively, as quickly as possible based on gut feelings via Chronos (Psychology Software Tools, Inc., Pittsburgh, USA). The mapping of valence to response keys was counterbalanced across participants. Crucially, participants were explicitly instructed to minimize the influence of their judgment of the preceding object. Participants completed this task twice: once preceding the cued recall phase and once following it.

#### Self-report Questionnaires

For insomnia measurements, participants completed the 7-item Insomnia Severity Index (ISI; Morin et al., 2011), which is a validated self-reported scale assessing insomnia severity across symptom domains. Participants rated the severity of symptoms over the preceding two weeks on a 5-point Likert scale (0 = Very Satisfied, 4 = Very Dissatisfied), with higher scores indicating more severe insomnia symptoms. Participants also completed the 21-item Beck Depression Inventory-II (BDI-Ⅱ; Beck, Steer, & Brown, 1996), which evaluated behavioral, cognitive, and affective symptoms of depression. Items were rated from 0 (symptom absent) to 4 (severe symptom intensity), with higher scores indicating more severe depression symptoms. For anxiety measurements, participants completed the 40-item Spielberger State-Trait Anxiety Inventory (STAI; Spielberger et al., 1971) that comprises two subscales: trait anxiety and state anxiety. Items were rated on a 4-point scale, with higher scores reflecting greater anxiety. Participants completed the Chinese version of these questionnaires, which were validated among the Chinese population (Chung, Kan, & Yeung, 2011; Huang et al., 2015; Li et al., 2021).

#### Statistical analysis

Facilitation (i.e., better memory for the Think condition compared to the Baseline condition) was assessed using the same model as the suppression-induced forgetting, except for the difference in the fixed effect of Condition (Think vs Baseline):

Identification or Gist or Detail ~ 1 + Group * Condition + age + depression + (1 | subject) + (1 | item)

Affective processing in the implicit affect task was quantified through positive classification probabilities per stimulus item. Cumulative link mixed models were performed with fixed effects of Group (healthy sleeper vs. insomnia), Condition (No-Think vs. Baseline), their interaction, and covariates (i.e., age and depression). Random intercepts for participants and items were incorporated:

Positive classification probabilities ~ 1 + Group * Condition + age + depression + (1 | subject) + (1 | item).

Two participants in the insomnia group were excluded from the implicit affect task preceding the cued recall phase due to pressing the same key during the whole task.

### Results

#### Suppression-induced forgetting: For Identification, the planned contrasts (Baseline vs. No-Think) based on group-level data showed a significant suppression-induced forgetting effect on Identification in healthy sleepers, *t*(39) = 4.20, *p* < .001, Cohen's *d* = 0.67, 95% CI [0.32, 1.00], but not in individuals with insomnia, *t*(39) = 1.55, *p* = .129, Cohen's *d* = 0.25, 95% CI [-0.07, 0.56]. For Gist, we observed a significant main effect of Condition (*F*(1, 1803.40) = 3.95, *p* = .047): Participants showed suppression-induced forgetting in Gist (*t* = 1.99, *p* = .047, Cohen’s *d* = 0.09, 95% CI [0.01, 0.18]). No significant Condition by Group interaction (*F*(1, 1804.37) = 0.03, *p* = .874) and main effect of Group (*F*(1, 76) = 0.28, *p* = .596) were observed. The marginal and conditional R^2^ of our model were 0.02 and 0.36, respectively. The planned contrasts (Baseline vs. No-Think) showed no significant suppression-induced forgetting effects for Gist in both groups (*p*s > .172). For Detail*,* we observed a significant Condition effect (*F*(1, 1803.25) = 14.67, *p* < .001; see Supplementary Figure 1b): Participants showed worse memories performance in the No-Think condition compared to the Baseline condition (*t*(1803) = -3.83, *p* < .001, Cohen’s *d* = -0.18, 95% CI [-0.27, -0.06]). There was no significant Condition by Group interaction (*F*(1, 1804.06) = 0.26, *p* = .609) and Group effect (*F*(1, 76.05) = 2.08, *p* = .153). The marginal and conditional R^2^ of our model were 0.03 and 0.43, respectively. The planned contrasts (Baseline vs. No-Think) revealed significant suppression-induced forgetting effects for Details in healthy sleepers, *t*(39) = 2.20, *p* = .034, Cohen's *d* = 0.37, 95% CI [0.03, 0.67], and individuals with insomnia, *t*(39) = 2.43, *p* = .020, Cohen's *d* = 0.38, 95% CI [0.06, 0.70].

#### Facilitation: For Identification, no significant main or interaction effects were found (*χ^2^*(1) < 1.90, *p*s > .168). The McFadden’s pseudo-R² for this model was low (pseudo-R² = 0.01). For Gist, we observed a significant Condition by Group interaction (*F*(1, 1804.30) = 4.64, *p* = .031; see Supplementary Figure 1): Healthy sleepers recalled more Gist in the Think condition compared to the Baseline condition (*M* = 4.9 %, *t*(1804) = 2.74, *p* = .006, Cohen’s *d* = 0.18, 95% CI [0.05, 0.31]), whereas participants with insomnia showed no significant facilitation (*M* < -0.2 %, *t*(1803) = -0.31, *p* = .756, Cohen’s *d* = -0.02, 95% CI [-0.15, 0.11]). Neither the main effect of the Group (*F*(1, 76) = 0.25, *p* = .618) nor Condition (*F*(1, 1803.20) = 2.94, *p* = .086) reached significance. The marginal and conditional R^2^ of our model were 0.02 and 0.34, respectively. For Detail, a significant Condition effect was observed (*F*(1, 1803.19) = 8.06, *p* = .005): Participant recall more Detail in the Think condition compared to the Baseline condition (*t*(1803) = 2.84, *p* = .005, Cohen’s *d* = 0.13, 95% CI [0.04, 0.22]). No main effect of Group (*F*(1, 76.04) = 0.87, *p* = .354) nor interaction effect (*F*(1, 1804.00) = 3.44, *p* = .064) was significant. The marginal and conditional R^2^ of our model were 0.03 and 0.42, respectively.

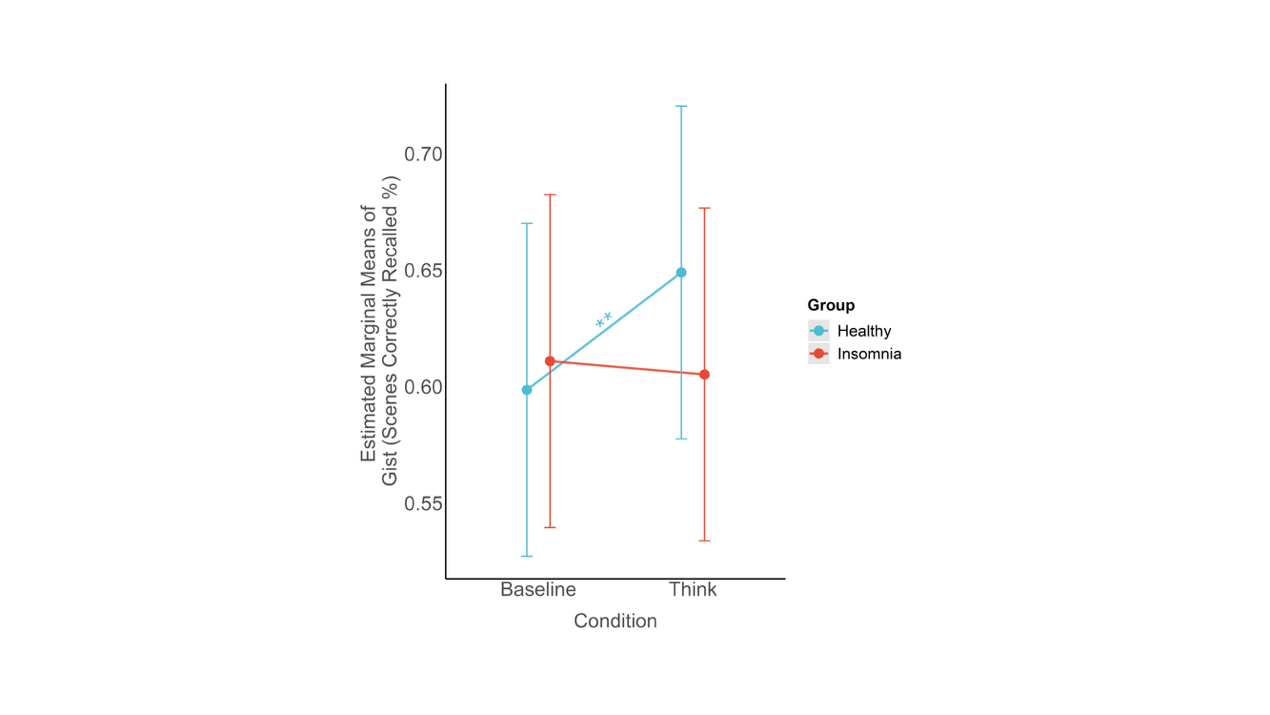

Supplementary Figure 1. Memory performance of Gist in the cued recall task: healthy sleepers showed better memories for the Think condition compared to the Baseline condition, suggesting a facilitation, whereas individuals with insomnia did not. Error bars represent ± 1 standard error. Asterisks present significant difference between conditions (^*^*p* < .05; ^**^*p* < .01; ^***^*p*< .001).

#### Neural activity during the Think/No-Think phase (Think vs. No-Think): For N450, we observed a significant main effect of Condition (*F*(1, 1826.92) = 8.85, *p* = .003): The No-Think condition elicited enhanced N450 amplitude compared to the Think condition (*t*(1827) = 2.97, *p* = .003, Cohen’s *d* = 0.14, 95% CI [0.05, 0.22]). No significant Condition by Group interaction (*F* (1,1828.13) = 0.01, *p* = .920) and Group effect (*F*(1, 76.99) = 0.38, *p* = .541) were observed. The marginal and conditional R^2^ for this model were 0.03 and 0.51, respectively. For P300, we observed a significant effect of Condition (*F*(1, 1827.99) = 14.54, *p* < .001): The Think condition elicited larger P300 amplitude compared to the No-Think condition (*t*(1827) = 3.81, *p* < .001, Cohen’s *d* = 0.17, 95% CI [0.08, 0.26]). No significant Condition by Group interaction (*F* (1, 1829.45) = 2.01, *p* = .157) and Group effect (*F* (1, 76.92) = 0.66, *p* = .420) were observed. The marginal and conditional R^2^ for this model were 0.01 and 0.40, respectively.

#### Implicit affect classification preceding the cued recall phase: We observed a significant Condition by Group interaction (*χ^2^*(1) = 6.91, *p* = .008; see Supplementary Figure 2a): Healthy sleepers showed significantly less positive classification probabilities for the No-Think condition compared to the Baseline condition (*M* = -4.2 %, *z* = -2.49, *p* = .013, OR = 0.74, 95% CI [0.58, 0.94]). In contrast, individuals with insomnia showed no significant difference between the No-think and Baseline conditions (*M* = 2 %, *z* = 1.21, *p* = .225, OR = 1.16, 95% CI [0.92, 1.46]). We found a significant Group effect (*χ^2^*(1) = 6.12, *p* = .013) and non-significant main effect of Condition (*χ^2^*(1) = 0.76, *p* = .384). The McFadden’s pseudo-R² for this model was low (pseudo-R² < .01).

Regarding the difference between the Think and the Baseline conditions, we found a significant Condition by Group interaction (*χ^2^*(1) = 4.45, *p* = .035; see Supplementary Figure 2b): Healthy sleepers showed significant less positive classification probabilities for the Think condition compared to the Baseline condition (*M* = -5.1 %, *z* = -2.34, *p* = .020, OR = 0.75, 95% CI [0.59, 0.96]), while individuals with insomnia showed no significant difference between the Think and the Baseline conditions (*M* = 1 %, *z* = 0.64, *p* = .523, OR = 1.08, 95% CI [0.85, 1.37]). We also found a significant Group effect (*χ^2^*(1) = 5.37, *p* = .020) and non-significant main effect of Condition (*χ^2^*(1) = 1.42, *p* = .233). The McFadden’s pseudo-R² for this model was low (pseudo-R² < .01).

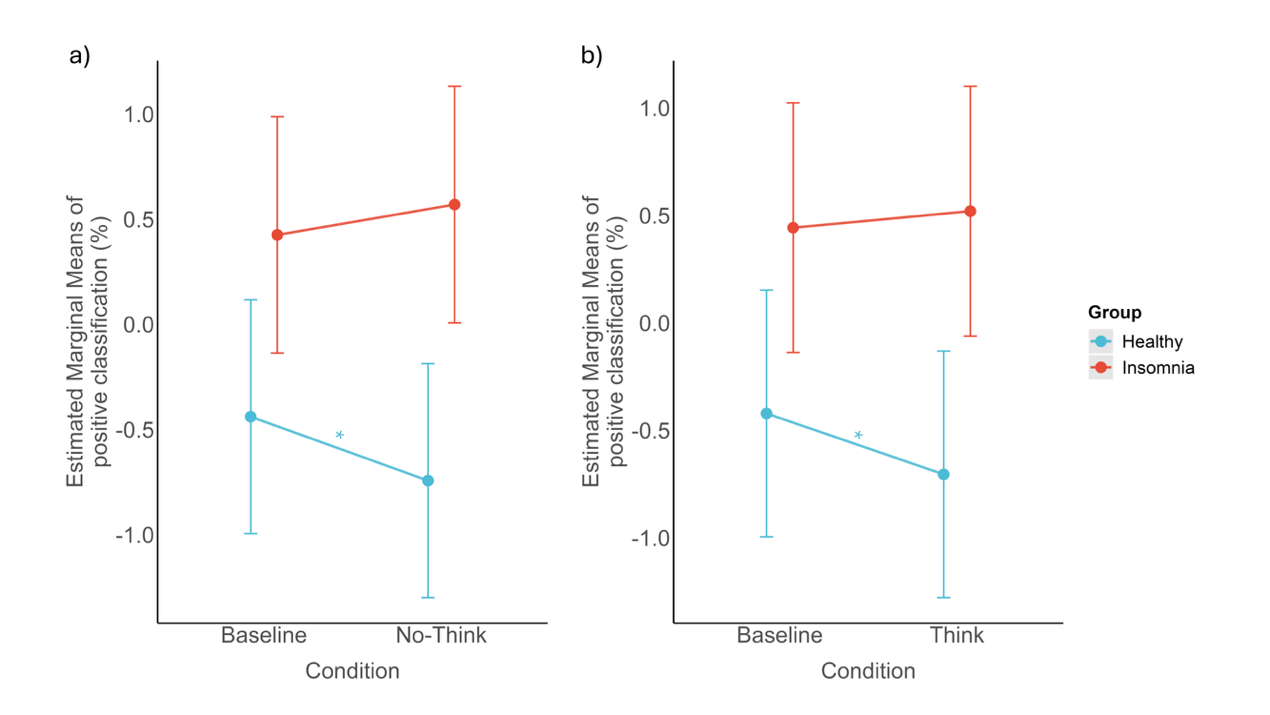

Supplementary Figure 2. Positive classification probabilities in the implicit affect task before the cued recall phase: healthy sleepers showed a) lower positive classification probabilities for the No-Think condition and b) lower positive classification probabilities for the Think condition, compared to the Baseline condition, respectively, whereas individuals with insomnia did not. Error bars represent ± 1 standard error. Asterisks present significant difference between conditions (^*^*p* < .05; ^**^*p* < .01; ^***^*p*< .001).

#### Implicit affect classification after the cued recall phase: Non-significant Condition by Group interaction was observed (*χ^2^*(1) = 0.37, *p* = .543). We observed a significant main effect of Condition (*χ^2^*(1) = 5.40, *p* = .020): Participants showed less positive classification probabilities for the No-Think condition compared to the Baseline condition (*z* = -2.34, *p* = .020, OR = 0.81, 95% CI [0.68, 0.97]). The Group effect was not significant (*χ^2^* (1) = 0.25, *p* = .617). Regarding the difference between Think and Baseline conditions, no significant main or interaction effects were found (*χ^2^*(1)s < 3.58, *p*s >.065). The McFadden’s pseudo-R² for this model was low (pseudo-R² < .01).

#### Neural activity during retrieval suppression (Think vs. No-Think vs. Perceptual Baseline): For N450, consistent with previous TNT research, the linear mixed-effects model revealed a significant Condition effect (*F*(2, 68.5) = 5.25, *p* = .007; see Supplementary Figure 3), with the Think condition eliciting significantly lower N450 amplitudes (i.e., more positive N450) than the No-Think condition (*t*(2306.1) = 2.97, *p* = .008, Cohen’s *d* = 0.13, 95% CI [0.05, 0.22]). Other comparisons were not significant (Perceptual Baseline vs No-Think, *p* = .178; Perceptual Baseline vs Think, *p* = .747). The main effect of Group (*F*(1, 77.4) = 0.20, *p* = .658) and the Condition by Group interaction was not significant (*F*(2, 2306.45) = 1.32, *p* = .267). The marginal and conditional R^2^ for this model were 0.03 and 0.50, respectively.

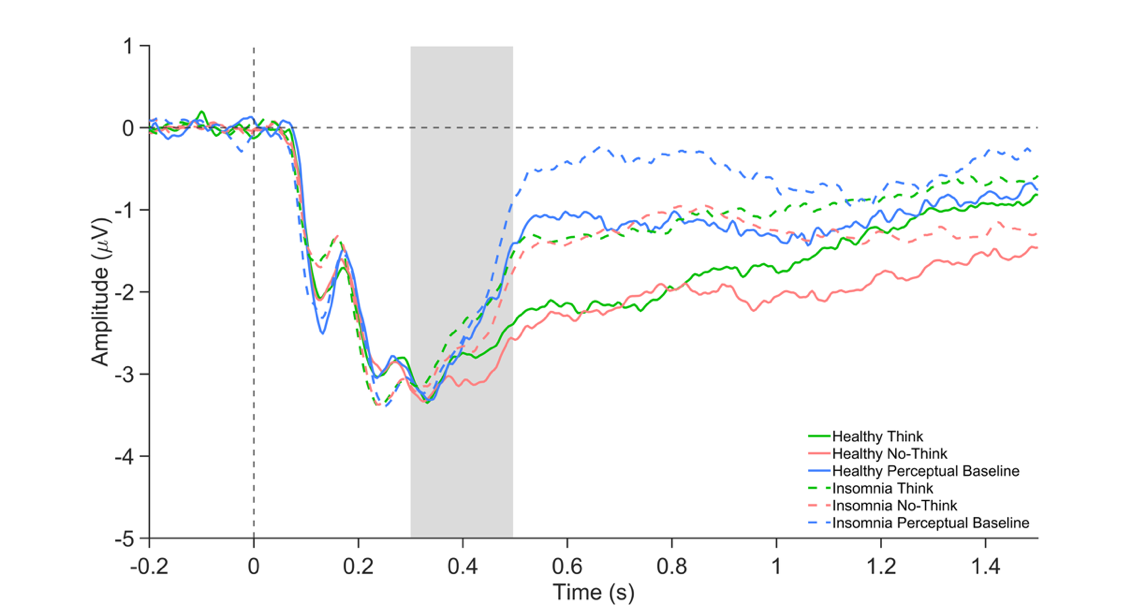

Supplementary Figure 3. The average ERP waveforms of the frontocentral (FC1, FC2, C1, C2, FCz, Cz) N450 from 300-500 ms in both groups.

For P300*,* a significant Condition by Group interaction was found (*F*(2, 2307.18) = 3.08, *p* = .046; see Supplementary Figure 4). Post-hoc analysis revealed that among healthy sleepers, the Think and the Perceptual Baseline conditions elicited larger P300 amplitude than the No-Think condition (Think vs. No-Think, *t*(2308.4) = 3.73, *p* = .001, Cohen’s *d* = 0.24, 95% CI [0.11, 0.37]; Perceptual baseline vs. No-Think, *t*(60.1) = 2.75, *p* = .021, Cohen’s *d* = 0.33, 95% CI [0.09, 0.58]). In contrast, participants with insomnia showed no significant P300 differences across conditions (*p*’s > 0.193). A significant Condition main effect was found (*F*(2, 74.18) = 8.08, *p* < .001), whereas the Group main effect was not significant (*F*(1, 77.53) = 0.27, *p* = .603). The marginal and conditional R^2^ for this model were 0.01 and 0.40, respectively.

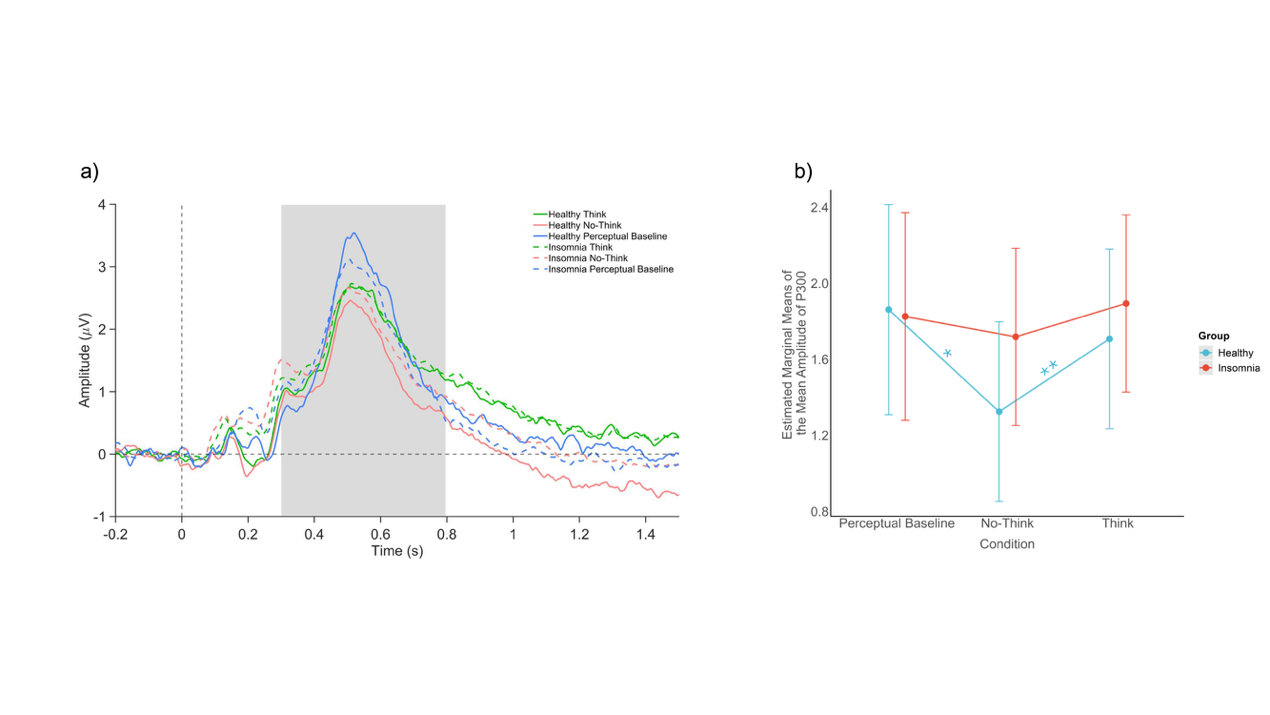

Supplementary Figure 4. a) The average ERP waveforms of the parietal (Pz, CP1, CP2, P1, P2) P300 from 300-800ms in both groups; b) Mean amplitude of P300 during the Think/No-Think phase: healthy sleepers showed larger P300 during the Perceptual Baseline condition and the Think condition, compared to the No-Think condition, whereas individuals with insomnia showed no such difference. Asterisks present significant difference between conditions (^*^*p* < .05; ^**^*p* < .01; ^***^*p*< .001).

For theta power, we found a significant Condition by Group interaction (*F* (2, 2307.37) = 6.17, *p* = .002; see Supplementary Figure 5): Healthy sleepers showed lower theta power during the Think condition relative to the No-Think condition (*t*(2311.1) = -3.52, *p* = .001, Cohen’s *d* = -0.23, 95% CI [-0.36, -0.10]), whereas participants with insomnia showed no significant differences between the Think and No-Think conditions (*t*(2306.8) = 0.01, *p* = .999, Cohen’s *d* < 0.01, 95% CI [-0.13, 0.13]). In addition, individuals with insomnia showed elevated theta power during the No-Think condition (*t*(85.4) = 2.41, *p* = .048, Cohen’s *d* = 0.22, 95% CI [0.04, 0.40]), and the Think condition (*t*(85.4) = 2.42, *p* = .046, Cohen’s *d* = 0.22, 95% CI [0.04, 0.40]), compared to the Perceptual Baseline condition, whereas healthy sleepers showed no significant differences between the Perceptual Baseline and the No-Think condition (*p* = .621), the Think condition (*p* = .284). Finally, individuals with insomnia showed larger theta power in the Think condition compared to healthy sleepers (*t*(88) = 2.38, *p* = .020, Cohen’s *d* = 0.47, 95% CI [0.08, 0.85]). A significant main effect of Condition was found (*F*(2, 53.93) = 4.08, *p* = .022, whereas the main effect of Group was not significant (*F*(1, 77.76) = 1.99, *p* = .163). The marginal and conditional R^2^ for this model were 0.04 and 0.34, respectively.

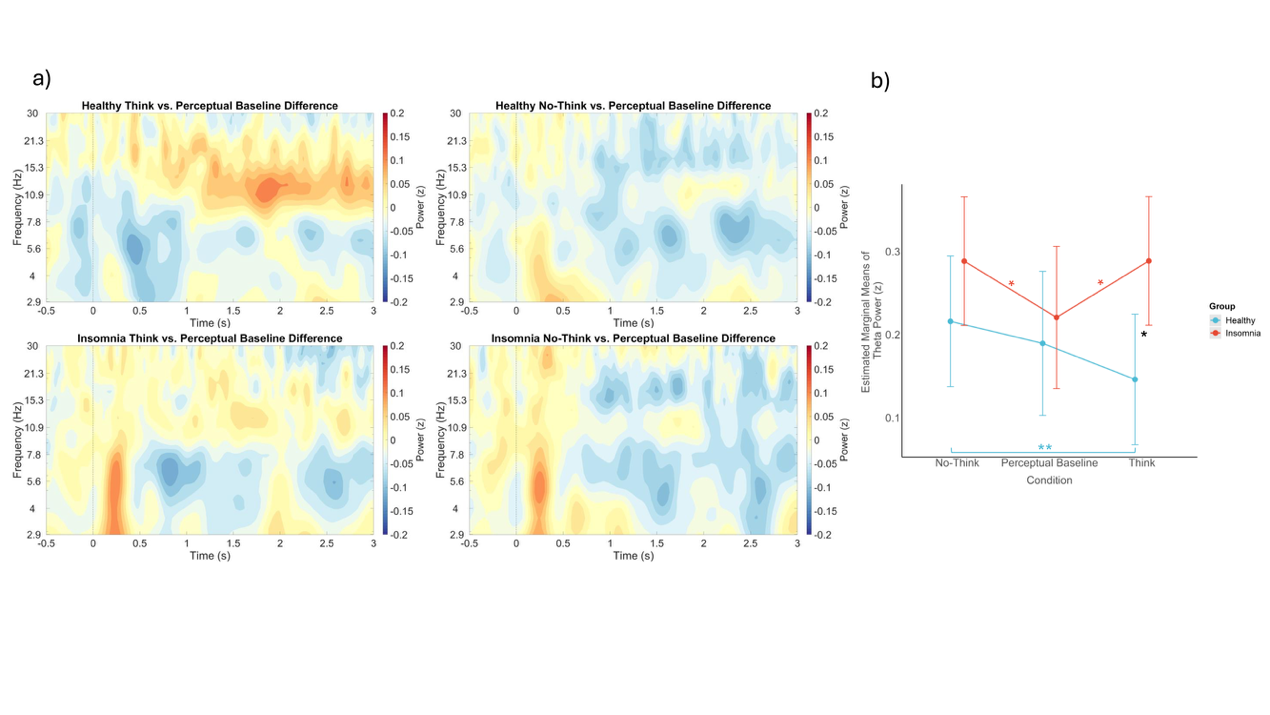

Supplementary Figure 5. a) The time-frequency plots of condition differences (subtracting the Perceptual Baseline condition from the Think/No-Think condition) in healthy sleepers and the insomnia group at the right prefrontal area (AF4, F4, F6). b) Theta power during the Think/No-Think phase: healthy sleepers exhibited enhanced right prefrontal theta power during the No-Think condition compared to the Think condition, whereas individuals with insomnia showed no such difference. Individuals with insomnia showed elevated theta power during the No-Think and Think conditions, compared to the Perceptual Baseline condition. Finally, individuals with insomnia exhibited enhanced theta power in the Think condition compared to healthy sleepers. Asterisks present significant difference between conditions (^*^*p* < .05; ^**^*p* < .01; ^***^*p*< .001).

#### Correlations between mental health measures, memory performance, and EEG effects among the insomnia group: A negative correlation was observed between the difference in right prefrontal theta power during the No-Think versus Perceptual condition and suppression-induced forgetting in Detail (rho = -0.31, *p* = .045, see Supplementary Figure 6). Specifically, higher suppression-induced forgetting effect in Detail was associated with enhanced theta power in the No-Think condition relative to the Perceptual Baseline condition.

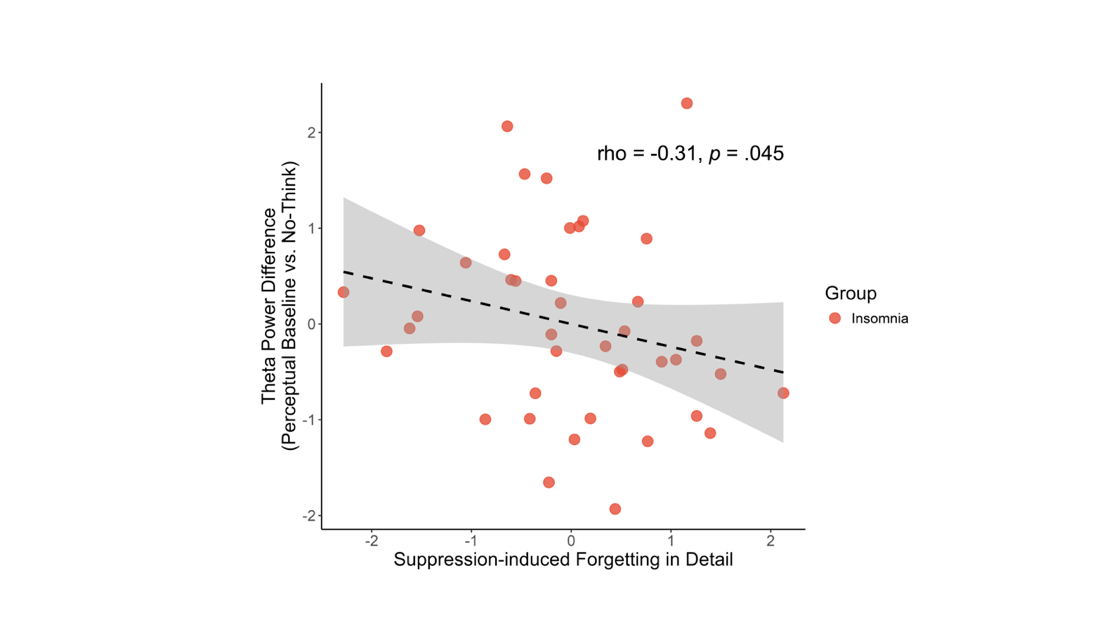

Supplementary Figure 6. Scatter plots depicted the correlation between z-normalized suppression-induced forgetting in Detail and theta power difference between the Perceptual Baseline and the No-Think conditions (subtracting the No-Think condition from the Perceptual Baseline condition) at the right prefrontal (AF4, F4, F6) area among individuals with insomnia.
